# Urinary F plasmids reduce permissivity to coliphage infection

**DOI:** 10.1101/2022.03.23.485578

**Authors:** Cesar Montelongo Hernandez, Catherine Putonti, Alan J. Wolfe

## Abstract

The microbial community of the urinary tract (urinary microbiota or urobiota) has been associated with human health. Bacteriophages (phages) and plasmids present in the urinary tract, like in other niches, may shape urinary bacteria dynamics. While urinary *E. coli*, often associated with urinary tract infection (UTI), and their phages have been catalogued for the urobiome, the dynamics of their interactions have yet to be explored. In this study, we characterized urinary *E. coli* F plasmids and their ability to decrease permissivity to *E. coli* phage (coliphage) infection. Putative F plasmids were present in 47 of 67 urinary *E. coli* isolates, and most of these plasmids carry genes that encode for toxin-antitoxin (TA) modules, antibiotic resistance, and/or virulence. Two urinary *E. coli* F plasmids, from urinary microbiota (UMB) 0928 and 1284, were conjugated into *E. coli* K-12 strains; plasmids included genes for antibiotic resistance and virulence. These plasmids, pU0928 and pU1284, decreased permissivity to coliphage infection by the laboratory phage P1vir and the urinary phages Greed and Lust. Furthermore, pU0928 could be maintained in *E. coli* K-12 for up to ten days in the absence of antibiotic resistance selection; this included maintenance of the antibiotic resistance phenotype and decreased permissivity to phage. Finally, we discuss how F plasmids present in urinary *E. coli* could play a role in coliphage dynamics and maintenance of antibiotic resistance in urinary *E. coli*.

**Importance:** The microbial community of the urinary tract (urinary microbiota or urobiota) has been associated with human health. Bacteriophages (phages) and plasmids present in the urinary tract, like in other niches, may shape urinary bacteria dynamics. While urinary *E. coli*, often associated with urinary tract infection, and their phages have been catalogued for the urobiome, the dynamics of their interactions have yet to be explored. In this study, we characterized urinary *E. coli* F plasmids and their ability to decrease permissivity to E. coli phage (coliphage) infection. Two urinary *E. coli* F plasmids, each encoding antibiotic resistance and transferred by conjugation into naïve laboratory *E. coli* K-12 strains decreased permissivity to coliphage infection. We propose a model by which F plasmids present in urinary *E. coli* could help to maintain antibiotic resistance of urinary *E. coli*.

## Background

The urinary tract is not sterile. It contains microbiota, including bacteria, eukaryotic viruses, fungi, archaea, and bacteriophages^1, 2^. The presence and proportion of specific bacteria in the urinary microbiota (urobiota) are linked to asymptomatic and symptomatic urinary conditions^1, 3^. Phages are key influencers of bacterial communities and, by extension, of human health^4–6^. Urinary phages are rich and diverse, both free-living and as prophages^7, 8^. As in other anatomical sites, urinary phage likely impact urobiota populations by influencing population structure, genetic exchange, and bacterial metabolism^4, 9–11^. In contrast, a bacterium may modulate a phage’s life cycle via traits encoded by its chromosome or by mobile genetic elements (MGE), such as plasmids^12–15^.

Plasmids can influence phage-bacteria dynamics by transmitting traits vertically and horizontally in bacterial populations^16–18^. For example, TraT can block both foreign plasmids and phage invasion^16, 19, 20^. Toxin-Antitoxin (TA) systems make plasmid loss and, by extension the antitoxin, lethal to the host; yet, TA modules can also antagonize phage life cycles^17, 21, 22^. In contrast, phage may target plasmid components, such as the conjugal apparatus plasmids use to transfer copies of themselves^23–25^. This can cause plasmid loss and result in antibiotic sensitivity, as many plasmids carry antibiotic resistance determinants^25–28^. Bacteria-plasmid-phage interactions have been primarily studied in laboratory settings and are yet to be thoroughly tested in complex communities. Bacteria-plasmid-phage interactions are multifaceted and could be key to understanding the dynamics and evolution of environmental microbial populations, such as the urobiota^13, 29, 30^. A key unanswered question is whether urinary plasmids and phage interact.

The best studied bacterium of the urinary tract is *E. coli*, often associated with urinary tract infection (UTI)^31, 32^. Malki et al. isolated seven *E. coli* phages (coliphages) from urine collected via catheterization from women with urge urinary incontinence^33^. Two of these coliphages, Greed and Lust, could infect and lyse *E. coli* strains^8^. Phage predation of *E. coli* in the urinary tract remains understudied, including the role plasmids play in urinary bacteria-phage dynamics^8, 34–37^. Previously, we described plasmids present in urinary *E. coli* isolates; most were F plasmids with genes for conjugation^38, 39^. F plasmids are genetically heterogenous and easily transmissible plasmids with high clinical relevance as they often carry genes encoding antibiotic resistance and virulence^40–42^.

We hypothesized that urinary *E. coli* F plasmids could influence permissivity to phage infection; we tested this hypothesis by monitoring lysis by coliphages P1vir, Greed, and Lust. P1vir is a laboratory phage mutated to only enter the lytic cycle; Greed and Lust are urinary phages that lyse *E. coli*, including some urinary strains^8, 33, 34, 43^. We found F plasmids in most of our urinary *E. coli* isolates; they often carried genes associated with TA modules, antibiotic resistance, and/or virulence. We then transferred 2 urinary F plasmids from their native hosts to laboratory *E. coli* K-12 strains; both reduced permissivity to infection by P1vir, Greed, and Lust. Both also transmitted antibiotic resistance and virulence genes. One, pU0928, was stable in *E. coli* K-12 for ten days without plasmid selection. We propose that F plasmids protect urinary *E. coli* from phage predation, maintaining antibiotic resistance in the population^44–46^.

## Methods

### Plasmidic assembly, genomic and gene homology scan, and annotation

We used urinary *E. coli* isolates and sequence read data previously published (BioProject PRJNA316969)^47^. Urinary isolates were previously recovered from urine samples obtained by transurethral catheterization from adult women during several IRB-approved studies at Loyola University Chicago (LU203986, LU205650, LU206449, LU206469, LU207102, and LU207152) and University of California San Diego (170077AW). Raw sequencing reads were assembled using plasmidspades.py of SPAdes v3.12 with k values of 55,77,99,127 and the only-assembler parameter^48, 49^. Plasmidic assemblies were queried against the nr/nt database via NCBI (web) BLAST to assign contigs as either plasmid or chromosome^50, 51^. Only contigs exhibiting sequence similarity to plasmids were examined. A homology heatmap of plasmidic assemblies was generated using sourmash v4.0 with parameters scaled=1000 and k=31^52^. For comparison, this included three urinary *Klebsiella pneumoniae* plasmidic assemblies, which were processed as described above.

We catalogued the plasmidic assemblies via PlasmidFinder v2.1 using the Enterobacteriaceae database with minimum thresholds of 95% identity and 60% minimum coverage^53^. Given their *inc* gene profile, we assigned plasmids as F plasmid (IncF group) or not (Non-IncF)^41, 54^. To identify known acquired antibiotic resistance genes, we scanned the urinary plasmidic assemblies with ResFinder v4.1 using the “acquired antimicrobial resistance genes” option and *Escherichia coli* species database^55^. To identify known virulence genes, the urinary plasmidic assemblies were scanned with VirulenceFinder v2.0 using the *Escherichia coli* species database, with identity threshold of 90% and minimum sequence length 60%^56^. Urinary plasmidic assemblies were annotated using Prokka v1.14.5 and sorted by length via the sortbyname.sh script from bbmap^57, 58^. Amino acid ORFs were clustered by homology using USEARCH v.11.0 with -cluster-fast -id 0.8 -clusters parameters^59^.

### Urinary plasmid conjugation and phenotype testing

Native antibiotic resistance cassettes were identified via ResFinder as described above. To validate predictions, urinary *E. coli* isolates were struck onto lysogeny broth (LB) plates supplemented with antibiotics: Ampicillin (Amp, 100 µg/ml), Chloramphenicol (Cam, 25 µg/ml), Kanamycin (Kan, 40 ug/ml), Spectinomycin (Spec, 100 µg/ml), or Tetracycline (Tet, 15 µg/ml). Both urinary and laboratory *E. coli* isolates were incubated overnight at 37°C.

To test urinary plasmid effects on phage infection permissiveness, conjugal plasmids from urinary *E. coli* isolates were transferred to derivatives of *E. coli* K-12 strain MG1655 using as described^39, 60–62^. These recipients included MG1655 transformed with multicopy plasmid pCA24n, encoding chloramphenicol resistance^61–63^. Other recipients were MG1655 with deletions of genes *yfiQ* and/or *cobB* marked by a resistance cassette. These genes are related to protein acetylation but exert no phenotype under growth conditions tested. pCA24n-*yfiQ* was purified from the ASKA collection^63^.

The lytic phages P1vir, Greed, and Lust were described previously^33, 34^. We titered them with the full plate titer technique^64^ and tested permissibility of *E. coli* isolates with the following phage spot titration assay. Briefly, *E. coli* transconjugants and controls were struck from frozen stocks onto selective plates and incubated overnight at 37°C. Single colonies were inoculated into 5 ml LB with appropriate selection and aerated overnight at 37°C. From each overnight culture, 100 ul was transferred to 5 ml of LB liquid with the appropriate antibiotic and aerated at 37°C until early exponential phase (OD_660_∼0.4, ∼3×10^8^ colony forming unit (CFU)/ml). From each subculture, 200 µl were transferred to a tube with 0.7% agar LB media preheated to 52°C, immediately mixed and plated onto an 1.5% agar LB plate. Plates cooled for 10 minutes and spotted with 10 µl of each phage as a 1:10 serial dilution in LB with a starting concentration of 10^10^ particle forming unit (PFU)/ml and a final concentration of 10^2^ PFU/ml. Phage spots dried for 20 minutes and plates incubated overnight at 37°C. The following day, the lowest titration that resulted in clearance was noted; an integer was given to each titration based on the number of dilutions it was removed from the starting concentration (the lowest titration being one dilution at 10^9^ PFU/ml and the highest being eight dilutions at 10^2^ PFU/ml). A control plate consisted of an MG1655-derivative lawn (200 µl of ∼3×10^8^ cfu/ml) spotted with P1vir, Greed, Lust (10 µl of 10^10^ PFU/ml) plus the negative controls (10 µl of LB; 10 µl of temperate phage Lambda).

To further assess the effect of urinary plasmids on phage permissiveness of the transconjugants, growth curves were performed. Transconjugants and controls were struck from frozen stocks on the appropriate antibiotic plate and incubated overnight at 37°C. Single colonies inoculated 10 ml LB with the appropriate antibiotic and aerated overnight at 37°C. From each overnight culture, 1 ml was transferred to 25 ml of LB liquid in a 250 ml flask to obtain approximately the same cell density (OD_660_ ∼0.2); subcultures were aerated at 37°C until early exponential phase (OD_660_ ∼0.4). Each phage (P1vir, Greed, Lust) was titrated and 0.5 ml added to the flask to achieve a multiplicity of infection (MOI) of 0.01 or 10.0, with a no phage control. Cultures were aerated at 37°C for 8 hours with OD_660_ measured every hour. Each treatment was repeated in triplicate.

To test antibiotic resistance after conjugation, the transconjugants (carrying pU0928, pU1284, or pU1223), their urinary plasmid donor isolate, and naïve recipient were grown on antibiotic plates (tetracycline, kanamycin, ampicillin, spectinomycin, chloramphenicol, LB control) as described above. To test plasmid stability, urinary isolate UMB0928, MG1655 pCA24, MG1655 pCA24 pU0928 and MG1655 *yfiQ*::cm were aerated at 37°C in 5ml of liquid LB in the absence of antibiotic selection for plasmid pU0928 for 10 days. Cultures were plated onto tetracycline (pU0928 selection marker) and LB plates in triplicate daily, incubated at 37°C overnight and colonies counted. Plasmid stability was calculated as the number of colonies on the tetracycline plate over the number colonies on the LB plate. A ratio of 1 indicates plasmid retention, while a ratio of 0 indicates plasmid loss. To assess phage permissivity after incubation in the absence of selection, on day 1 and 10, the isolates were grown overnight to perform a spot titration assay and struck onto antibiotic plates, as described above.

### Urinary plasmid extraction, sequencing, and analysis

We extracted and sequenced the genomes from the transconjugants as described previously ^38^. Plasmid analysis of the transconjugants was performed as described above. To assess gaps in plasmid sequence coverage, raw sequencing reads were mapped to the curated plasmid assemblies via the BBmap plugin in Geneious^65^. The urinary plasmid sequences from the transconjugants (pU0928, pU1223, and pU1284) were scanned for phage genetic content via PHAST and PHASTER using default settings^66, 67^. Phage-like sequences predicted by PHAST and PHASTER were aligned to assess redundant sequences, which were pruned. The phage-like sequences from pU0928, pU1223, and pU1284 were compared to one another on overall sequence homology via sourmash^52^. The phage-like sequences from pU0928, pU1223, and pU1284 were annotated using Prokka v1.14.5 and processed as outlined above^57^. ORFs in plasmids pU0928, pU1223, and pU1284 were clustered by homology using USEARCH v.11.0 with - cluster-fast -id 0.8 -clusters parameters^59^

### Data Availability

Sequences will be made publicly available prior to publication.

## Results

### Urinary *E. coli* plasmid sequence analysis

From the 67 urinary *E. coli* genomic raw sequence reads, 57 contained putative plasmid sequences (**Supplemental Table 1**). We aimed to identify urinary *E. coli* isolates likely to carry F plasmids (even if the host carried other plasmids); therefore, plasmidic assemblies in each isolate were treated as a singular plasmidic unit and analyzed for F plasmid content. All manually curated plasmidic assemblies had homology to plasmid entries in the NCBI database (query coverage 71-100% with sequence identity 96-100%) (**Supplemental Table 1**). Homology of plasmidic assemblies was primarily to *E. coli* plasmid entries but also to plasmids from other Enterobacteriaceae, including *Klebsiella*, *Shigella*, and *Enterobacter*. Two plasmid assemblies shared homology with the plasmid in UPEC strain UTI89.

Plasmid assemblies were scanned for plasmid incompatibility genes (**Supplemental Table 2**). Fifty-seven urinary *E. coli* isolates were predicted to carry *inc* genes organized into two groups: either containing at least one *incF* gene (IncF group, n=47) or no *incF* genes (Non-IncF group, n=10) (**Supplemental Table 2**). While non-*incF* genes were identified in plasmidic assemblies, the most common incompatibility genes were from *incF*, which are associated with F plasmids (present in 68.7% of all profiled urinary *E. coli* strains and in 82.5% of those predicted to carry plasmids). We compared the overall sequence homology of the urinary plasmidic assemblies; the IncF group clustered into multiple subgroups. Plasmidic assemblies of the incF group were predicted to possess a total of 2,060 unique ORFs, while the Non-IncF group had a total of 895 unique ORFs. Only ∼24% of all plasmid ORFs were assigned a known function. The incF group had the highest count of distinct ORFs with assigned function, including annotations for plasmid replication machinery, metal transport and resistance genes, leukotoxin genes, multi-drug transporters, phage genes, and virulence regulators (data not shown).

Plasmidic assemblies were profiled for TA, antibiotic resistance, and virulence genes. Sixteen TA genes were predicted via Prokka annotation in the IncF group plasmidic assemblies but none in the Non-IncF group. Complete TA pairs were identified for *ccdAB, isoAB, mazEF, parDE,* and *pemIK*; *ccdAB* and *pemIK* were the most frequent with some plasmidic assemblies having hits for both modules (**Figure 1a**). The plasmidic assemblies were predicted to confer resistance to the following antibiotics: aminoglycoside, fluoroquinolone, macrolide, streptomycin, sulfonamide, tetracycline, and trimethoprim (**Figure 1b**). Some plasmidic assemblies were predicted to have no antibiotic resistance genes; in contrast, four F plasmidic assemblies (from UMB0906, UMB0949, UMB3538, UMB5924) were predicted to have seven antibiotic resistance genes (**Supplemental Figure 1a**). Many of the strains were able to grow on one or more antibiotics (**Supplemental Figure 1 b, 1c**). Thirty distinct virulence genes were predicted in the plasmidic assemblies (**Figure 1c**), with *traT* (78.72%) and *senB* (53.19%) the most common in the incF group. Non-IncF plasmidic assemblies had hits primarily to colicin-related virulence genes (*ccl, celb, cib, cia*), which are signature genes of Colicin plasmids, although the plasmidic assembly of the Non-IncF plasmidic assembly from UMB0731 had hits to *traT* and *senB*.

**Figure 1.**
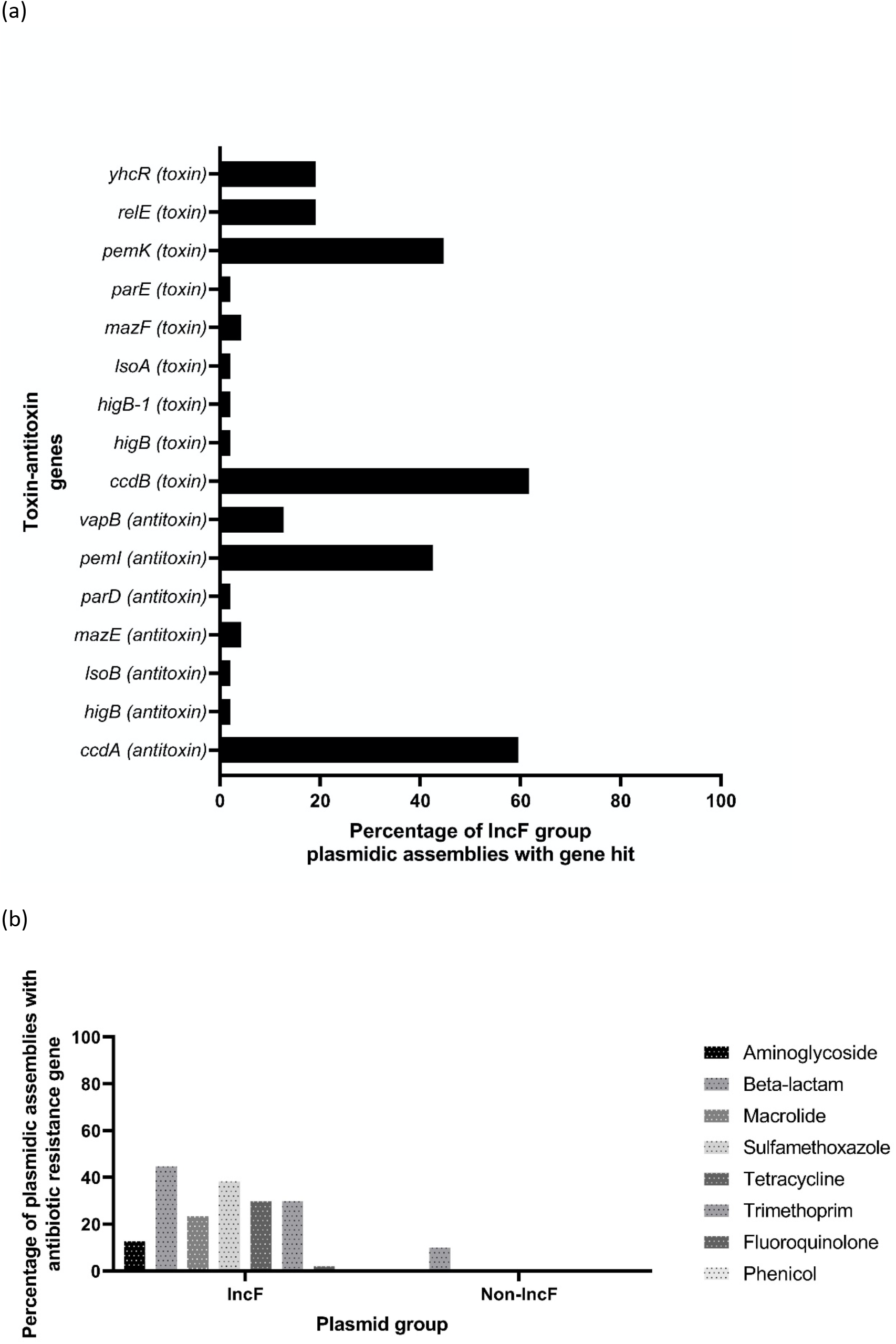
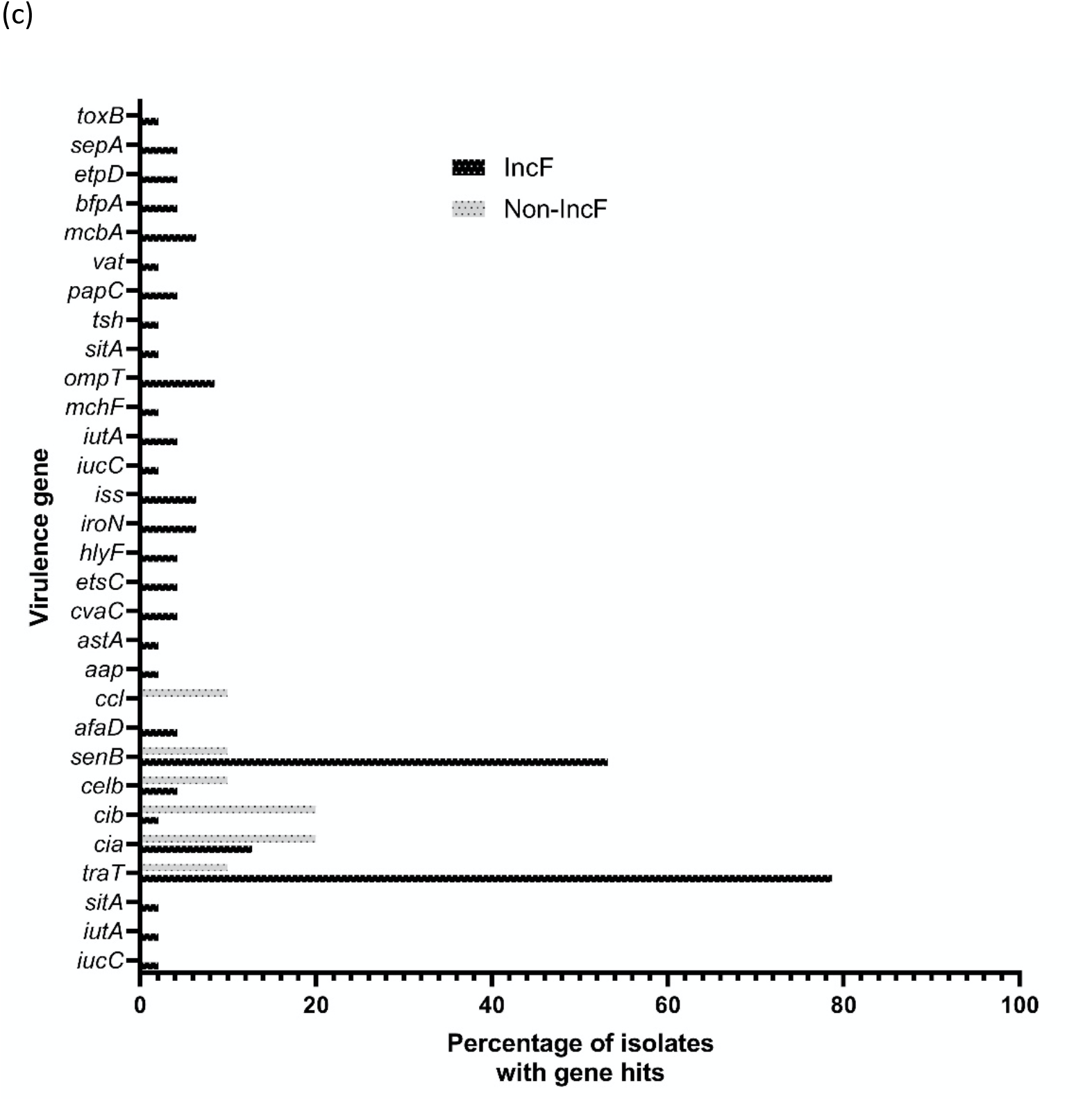
Proportion of plasmid addiction, antibiotic resistance, and virulence genes in urinary E. coli plasmidic assemblies (IncF=47, Non-IncF=11 plasmidic assemblies). Percentages denote plasmidic assembly Inc group with gene over that Inc group’s plasmidic assembly total. (a) Urinary E. coli F plasmid assemblies have a variety of TA genes which are associated with plasmid retention. ccdAB and pemIK are the most common modules. TA genes were not found in Non-IncF plasmidic assemblies. (b) Types of antibiotic resistances predicted in plasmid types in urinary E. coli. IncF group plasmidic assemblies have a higher proportion of antibiotic resistance genes compared to non-IncF group. (c) Percentage of isolates from each plasmid group predicted to have a given virulence gene. IncF group plasmidic assemblies had the largest variety and proportion of virulence gene hits. The most common virulence genes were traT (blocks invading plasmids) and senB (F plasmid-linked enterotoxin).

### Urinary plasmid transconjugant phenotype testing

To obtain a visual reference for future assays, we spotted a lawn of *E. coli* K-12 (derivatives of strain MG1655: MG1655 pCA24n-cm and MG1655 *ΔcobB yfiQ*::cm) with the lytic phages P1vir, Greed, and Lust; for controls, we used the temperate phage Lambda and LB. P1vir, Greed, and Lust resulted in cleared zones, while Lambda caused a turbid phenotype, and LB had no effect. We conjugated putative F plasmids from urinary *E. coli* (UMB0928, UMB1091, UMB1223, UMB1284, UMB6721) to multiple MG1655-derived strains, as described previously^39^. We then tested the susceptibilities of urinary *E. coli* and the transconjugants. The urinary plasmid donors were not permissive to the phage at any of the titers tested (**Supplemental Table 3**). In contrast, the naive recipients were susceptible at every titer tested, including dilution by eight orders of magnitude to MOI ∼10^-6^ (10^2^ PFU/mL, 10^8^ CFU/ml). When exposed to phage, phenotypes of transconjugants with either pU1091, pU1223, or pU6721 resembled those of naïve parent. In contrast, transconjugants with either pU0928 or pU1284 were susceptible only at the highest titers of P1vir, Greed, and Lust, but no observable clearing after 3-4 phage concentration titrations (i.e., decreased permissivity to infection). Multiple MG1655 derivatives were tested as recipients, with results consistent in all recipients tested. Given that pU0928 and pU1284 changed the spot titration phenotype, we further tested these plasmids using pU1223 as the negative control.

Growth curves assessed the effect of the urinary plasmids on *E. coli* growth during phage infection at MOIs of 0, 0.01, and 10.0 (**Figure2a-c**). P1vir Infection of transconjugants with pU0928 and pU1284 but not pU1223 resulted in comparable optical density to controls uninfected with phage at all time points (**Figure 2a, Supplemental Figure 2a**). Greed infection at MOI 0.01 of pU0928 and pU1284 but not pU1223 transconjugants resulted in growth like the uninfected control (**Figure 2b**, **Supplemental Figure 2b**). Increasing the MOI of Greed to 10.0 resulted in growth comparable to the recipient parents without pU0928 or pU1284 infected at MOI 0.01. Lust infection of the pU0928 and pU1284 transconjugants gave results like P1vir at the MOI tested (**Figure 2c**, **Supplemental Figure 2c**). In contrast, the pU1223 transconjugants had results comparable to the naïve recipients, indicating that pU1223 does not change growth (**Figure 2a-c, Supplemental Figure 2a-c**).

**Figure 2.**
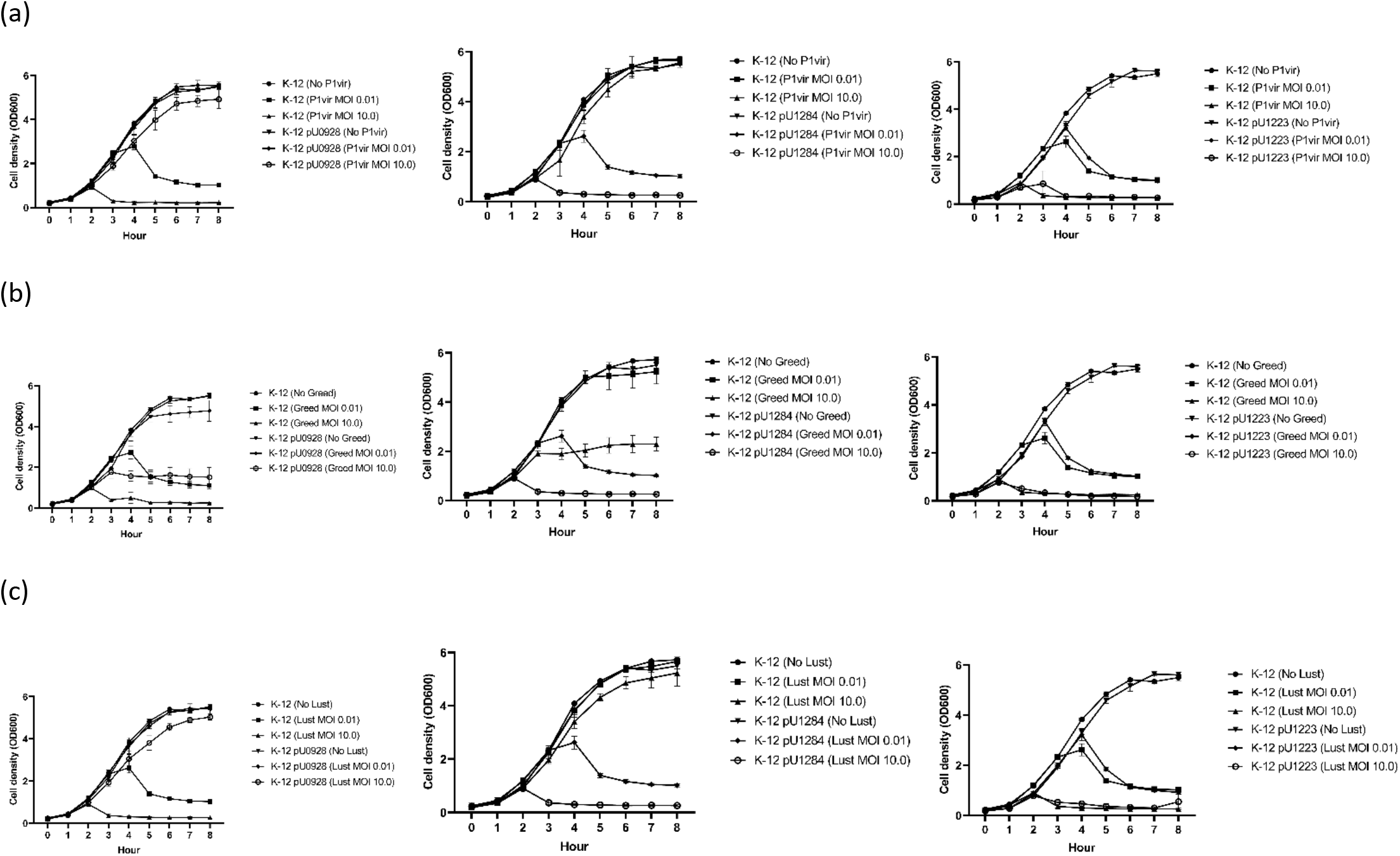
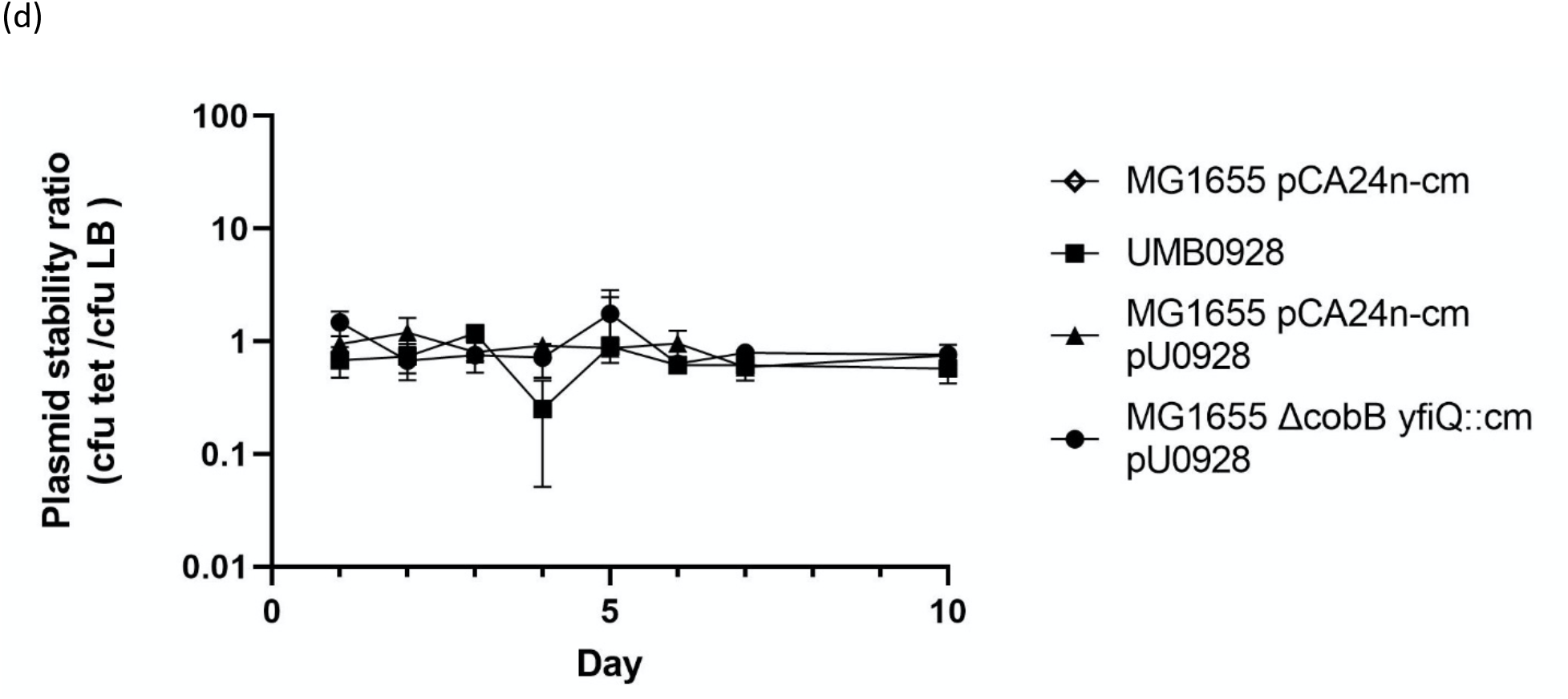
Growth of *E. coli* K-12 infected with phage and stability of urinary plasmids. Transconjugants *E. coli* K-12 (MG1655 pCA24n-cm) with pU0928, pU1284, or pU1223 infected with phage (0.01, 10.0) (a) P1vir, (b) Greed, and (c) Lust. Growth of naïve K-12 decreases with MOI of 0.01 and 10.0. pU0928 and pU1284 decreased permissivity to phage infection but pU1223 did not. Similar results observed when a different K-12 strain (MG1655 *ΔcobB yfiQ::Cm*) was used, see Supplemental Figures 4a-c. (d) Urinary isolate UMB0928 and *E. coli* K-12 variants were grown in the absence of antibiotic selection for plasmid pU0928 for 10 days. Cultures were plated onto tetracycline (pU0928 selection marker) and LB plates daily. A plasmid stability ratio (CFU in tetracycline divided by CFU in LB) of 1 indicates plasmid retention, while a ratio close to 0 indicates loss of plasmid. The negative control MG1655 pCA24n-cm without pU0928 did not grow on tetracycline plates.

Transconjugants exhibited growth on antibiotic plates like that of the respective donor parent (**Supplemental Table 4**). The stability of pU0928 was tested by incubation for multiple days in LB without the antibiotic that selects for the plasmid (i.e., tetracycline). After 10 days, growth on LB plates was comparable to growth on LB + tetracycline plates (**Figure 2d**). Overnight cultures from days one and ten were used to grow the bacteria on antibiotic plates and do a phage spot titration assay. Growth on antibiotic plates and phage permissivity profiles did not differ between day one and day 10 (**Table 1**).

**Table 1.**
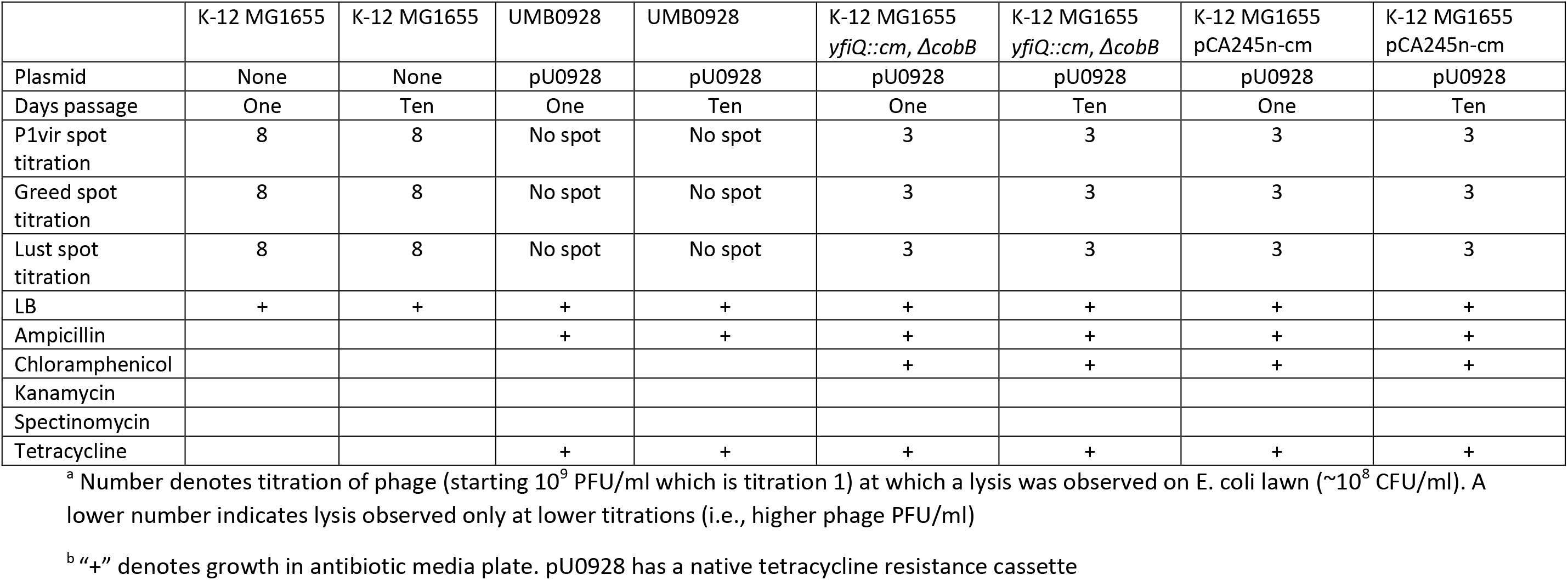
Phage spot titration and antibiotic plate growth phenotype (overnight compared to incubation for 10-days)**^a,b^**.

### Analysis of urinary plasmids in *E. coli* K-12 transconjugant

Of the urinary plasmids tested thus far, pU0928 and pU1284 reduced permissivity to phage infection; in contrast, pU1223 was one of the plasmids that did not change the infection phenotype relative to naïve recipients. We sequenced the transconjugants and explored sequences of the plasmids pU0928, pU1223, and pU1284; each was predicted to be ∼100k bp (approximately the size of a typical F plasmid), each had F plasmid *inc* genes, and each had sequence similarity (>45% query coverage and >99% percent identity) to F plasmid entries in the NCBI database (**Table 2**). Plasmids pU0928, pU1223, and pU1284 were predicted to have multiple antibiotic resistance genes, including tetracycline, which had been used as a selection marker (**Table 2**). No large gaps were observed in the plasmid sequence after mapping reads to the predicted plasmid sequence. Known gene functions were assigned to 39.05% of ORFs in pU0928, 40.82% of ORFs in pU1284, and 42.18% of ORFs in pU1223. The names of distinct ORFs were processed as a list for each plasmid. The plasmid sequences had ORFs annotated with plasmid replication, conjugation, maintenance machinery, and virulence functions.

**Table 2.**
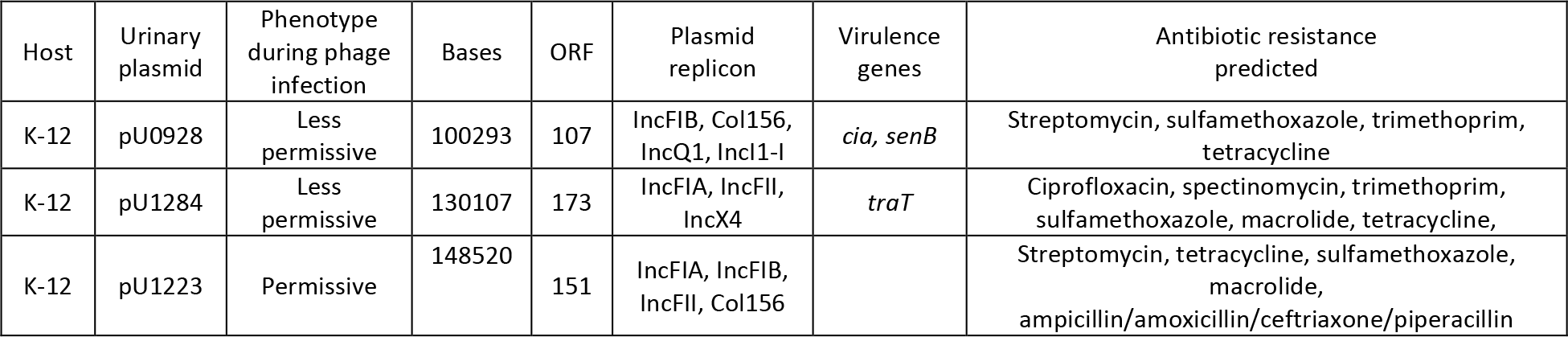
Overview of urinary plasmids conjugated into *E. coli* K-12.

The anti-phage F plasmids pU0928 and pU1284 were reviewed for genes that may antagonize phage infection and thus explain the change in phage infection phenotype (**Table 3**). Plasmid pU0928 is predicted to carry the immunity (*imm*) gene characterized in phage T4 and involved in phage superinfection exclusion^68, 69^. pU1284 is predicted to possess the gene *traT* reported to block phage adsorption, although the phage-permissive plasmid pU1223 also had this gene^16, 19, 20^. All three plasmids carry the TA modules *pemIK* and *ccdAB*. However, we identified three predicted ORFs in phage-nonpermissive plasmids pU0928 and pU1284 but not in the phage-permissive plasmid pU1223 sequence. We predicted these ORFs encode a phage integrase, a dihydrofolate reductase enzyme, and an EAL cyclic di-GMP phosphodiesterase domain-containing protein. These ORFs were highly conserved (>99% identity) between pU0928 and pU1284. The phage integrase ORF was highly conserved in 18 of the urinary *E. coli* plasmidic assemblies, including the UMB0928 and UMB1284 ancestral strains (**Supplemental Table 5**). All plasmid assemblies with this phage integrase were in the IncF group (38.3% of IncF plasmidic assemblies); the majority had low permissivity to phage infection, and presence of the phage integrase often co-occurred with the predicted dihydrofolate reductase or EAL cyclic di-GMP phosphodiesterase domain-containing protein.

**Table 3.** Coding regions in urinary plasmids with functions linked to phage infection^a^.

|  |  |  | pU0928 | pU1284 | pU1223 |
| --- | --- | --- | --- | --- | --- |
| Predicted coding region | Predicted gene function | Uniprot entry | K-12 less permissive to phage with plasmid | K-12 less permissive to phage with plasmid | K-12 permissive to phage just like WT |
| CcdA | Toxin of ccdAB module | P62552 | + | + | + |
| CcdB | Antitoxin of ccdAB module | P62554 | + | + | + |
| Dihydrofolate reductase | Phage DNA synthesis and protein assembly | P27422 | + | + |  |
| Imm | Phage superinfection exclusion | Q03708 | + |  |  |
| PemI | Antitoxin of pemIK module | P13975 | + | + | + |
| PemK | Toxin of pemIK module | P13976 | + | + | + |
| Phage integrase | Integrates phage genetic material into host genome | P62590 | + | + |  |
| TraT | Blocks phage adsorption | B1VCB1 | + | + | + |

<sup>a</sup>“+” denotes presence of amino acid sequence cover query and identity of over 90%.

Given that phage integrases are associated with prophage, the plasmids pU0928, pU1223, and pU1284 were scanned for other phage-like sequences (**Supplemental Table 6**). There were four distinct putative phage-like sequences present in pU0928, three in pU1223, and one in pU1284 (**Supplemental Figure 3**). The phage-like sequence regions pU0928_phaster_1, pU1223_phaster_1, pU1223_phast_3, and pU1284_phast_1 had a score of >90, indicating high likelihood of being an intact phage. The anti-phage pU0928 and phage-permissive pU1223 shared a predicted phage sequence (pU0928_phaster_1 and pU1223_phaster_1). The anti-phage pU0928 and pU1223 did not share phage-like sequences. Four ORFs were shared between pU0928_phast_1 and pU1284_phast_1, including two transposases, dihydrofolate reductase, and a phage integrase. The dihydrofolate reductase and phage integrase ORFs were the same identified as shared by the plasmids pU0928 and pU1284. The urinary plasmids previously predicted to contain this phage integrase were scanned for phage content via PHASTER. Except for UMB3641, all urinary plasmids were predicted to contain at least one phage-like sequence, with varying degrees of completeness, including those predicted to contain an intact phage (>90 score) (**Supplemental Table 7**).

## Discussion

Plasmids and phages are MGEs that impose distinct selective pressure on bacteria^11, 30, 44, 46^. We understand more of the complexity of plasmid and phage dynamics, but less of the role that plasmids may play in phage predation of *E. coli*, such as strains present in the urobiota^13, 29, 46^. *E. coli* is the urinary bacterial species most associated with UTI; its management can be difficult due to virulence factors and antibiotic resistance, often encoded by plasmids^31, 41, 70, 71^. F plasmids are easily transmissible, persistent, and often carry antibiotic resistance and virulence traits^19, 42, 72^. Despite *E. coli* being the most studied of bacterial species, only ∼25% of plasmid ORFs in urinary F plasmids were annotated with a known function ^38, 72–74^. In characterizing urinary *E. coli* plasmids, we took special attention in profiling genes involved in plasmid retention (i.e., TAs), antibiotic resistance, and virulence^41^.

IncF group plasmidic assemblies were predicted to have multiple antibiotic resistance genes and grew on multiple types of antibiotics (**Figure 1b**, **Supplemental Figure 1a-c**). In a previous study, we catalogued conjugation systems in urinary *E. coli* isolates, and we used this information to transfer F plasmids from urinary strains (UMB0928, UMB1223, and UMB1284) to *E. coli* K-12 strain MG1655 derivatives (**Supplemental Table 4**)^39^. Rather than modify the urinary plasmids, native antibiotic resistance on urinary plasmids was exploited as a selection marker during experiments. For pU0928, its multiple antibiotic resistances were stably maintained in MG1655 derivatives for ten days even in the absence of selection (i.e., passaged on LB), potentially explained by the TA modules *pemIK* and *ccdAB* (**Figure 2d**)^22, 75, 76^. TA modules, the most common being *mazEF* and *pemIK*, were present in urinary F plasmids (**Figure 1a**). Virulence genes were also present in the IncF group in a higher proportion and diversity relative to the Non-IncF group (**Figure 1c**). After conjugation of urinary plasmids into an MG1655 derivative, multiple virulence genes were now detected in the transconjugants. Taken together, these data show that urinary F plasmids not only code for antibiotic resistance and virulence genes, but also that these genes can be transferred to a naïve strain and stably maintained potentially due TA modules, a relevant factor for *E. coli* populations in the urinary tract^22, 77^. Following profiling urinary F plasmids, the key unanswered question was how these interact with phage.

We showed evidence that two urinary F plasmids, pU0928 and pU1284, could decrease their host’s permissivity to phage infection (**Figure 2a-c, Supplemental Table 3, Supplemental Figure 2**) but not provide immunity, as a high titer of phage (e.g., 10^10^ PFU/ml of P1vir, MOI 10^2^ PFU/ml) could still result in lysis, but not at lower titers. In contrast, the urinary *E. coli* used as plasmid donors were not lysed even at the highest concentration of phage tested. This plasmid-borne protective effect was observable not just in overnight transconjugant cultures but present even after passaging transconjugants for ten days in the absence of plasmid selection (**Table 1**). These results included testing with the lytic urinary phage Greed and Lust. While not thoroughly studied, phage titers in natural environments have been estimated to be in 10^7^ PFU/ml in soil samples and range from 10^2^-10^5^ PFU/ml in marine samples^78–80^. Therefore, we can infer that in low biomass environments, such as the urinary tract, these anti-phage plasmids could provide an edge to *E. coli* under phage predation^8^. Phage predation could be a selective force for phage-protective plasmids in urinary bacteria^13, 14, 45^. This scenario is akin to antibiotic utilization in the presence of bacteria carrying resistance plasmids: bacteria with plasmid-borne antibiotic resistance will survive, propagate, and thus increase the frequency of the plasmid^26, 44, 81^.

That stated, the mechanism by which the F plasmids protect urinary *E. coli* from phage predation remains unknown. Given our results, we know that traits expressed by the urinary plasmid did not confer immunity (i.e., zero permissivity to infection), but rather provides protection below a given MOI. Rather than plasmids granting infection immunity, akin to a phage receptor mutation, our data supports a stoichiometric relationship between the plasmid’s mechanism and infecting phage particles (**Figure 2a-c, Supplemental Table 3, Supplemental Figure 2**). In their study of bacteria-plasmid-phage interactions, Harrison et al. posit that conditions that limit extinction may stabilize phage-bacteria co-existence^46^. In this scenario, a mechanism that reduces permissivity to infection may be more stable in the long term than one that attempts infection immunity^29^. In terms of mechanism, we must highlight that phage permissivity was relatively comparable whether transconjugants were infected with P1vir, Greed, or Lust. P1vir is a temperate *Myovirida*e phage (genus: Punavirus) modified to only undergo the lytic life cycle; its structure consists of an icosahedral head connected to a tail with six tail fibers^34, 82, 83^. Greed and Lust are from the *Siphoviridae* family (genus: Seuratvirus) and related to the phages Seurat and CAjan, but still genetically distinct from each other; both phages were noted on transmission electron microscopy to have a head connected with a tail, with tail fibers predicted in their genome^33, 34, 84^. We hypothesize that the mechanism that decreases infection permissivity does not target a factor exclusive to a single phage, but potentially a conserved mechanism in the three phages, perhaps related to adsorption or the lytic pathway^14, 85, 86^.

To better understand pU0928 and pU1284, we extracted the genetic content in the transconjugants and analyzed the plasmid sequence (in addition to permissive pU1223). The three urinary plasmids were dissimilar in overall sequence, outside of being F plasmids, and only ∼25% of ORFs were assigned a function by Prokka. In terms of known anti-phage genes, all three plasmids had the TA modules *pemIK* and *ccdAB*, although only pU0928 had the superinfection exclusion gene *imm*, and only pU1223 and pU1284 had the phage adsorption blocking gene *traT* (**Table 3**)^16, 19, 22, 68, 69^. Given that pU0928 and pU1284 had similar protective phenotypes during phage infection, we hypothesize they could share similar ORFs linked to this mechanism. Three ORFs were present in pU0928 and pU1284 but absent from the permissive pU1223: a phage integrase, the enzyme dihydrofolate reductase, and an EAL cyclic di-GMP phosphodiesterase domain-containing protein. The former two genes can be linked to phage biology^87–90^.

Certain phages, such as Lambda, utilize phage integrases to integrate their genome into the host genome^35, 91^. A phage integrase is a signature for prophage, though by itself does not indicate viability^88, 92, 93^. Dihydrofolate reductase reduces dihydrofolic acid to tetrahydrofolic acid, involved in the amino and nucleic acid synthesis^89^. In phage, this enzyme plays a role in DNA synthesis, but has also been linked to proper packaging of the capsid; the enzyme must be finely regulated to achieve proper competition of the phage life cycle^89, 90^. This enzyme can be crucial such that some phages code their own dihydrofolate reductase, which replaces the host enzyme during the infection process^90^. Potentially, the dihydrofolate reductase in the plasmid could compete or otherwise interfere with the propagation of invading phage. The role of the EAL cyclic di-GMP phosphodiesterase domain-containing protein is more nebulous, but could be important to signaling^94^. Despite the overall low homology of ∼14% between the plasmids pU0928 and pU1284, the three shared ORFs were highly conserved at the amino acid level (the phage integrase was 99% identical, while the other two sequences were identical).

Given that phage integrases are associated with prophage, we scanned pU0928, pU1284 and pU1223 for phage sequences. Both pU0928 and pU1284 had predicted phage sequences, including some with high degree of sequence similarity to known phages (**Supplemental Table 6**). The low homology of the phage-like sequences in pU0928 (pU0928_phast_4) and pU1284 (pU1284_phast_1) indicates these sequences are distinct (**Supplemental Figure 3**), but these sequences do share some ORFs, including the phage integrase and dihydrofolate reductase described for the overall plasmid sequence. The anti-phage phenotype in pU0928 and pU1284 could be explained by the presence of distinct prophages that nonetheless share a protective mechanism^95^. It must be noted that only pU1284_phast_1 had a score >90 predicting it to be an intact phage, while pU0928_phast_4 had a score of 40, which is well below the confidence threshold. Alternatively, the lower score could indicate that plasmids have acquired phage-like genes or that these are remnants of past phage integrations, and thus not include a viable phage^96, 97^. There are reports of prophage integrating into plasmids, and prophage superinfection immunity and exclusion have been documented, but to our knowledge there are no reports of F plasmids protecting *E. coli* from phage infection^68, 95, 98^. Future studies should focus on identifying genes responsible for the anti-phage mechanism in F plasmids. Fortunately, this may be a realistic endeavor given that pU0928 and pU1284 are now in the genetically tractable *E. coli* K-12 MG1655^99^.

### Phage and urinary *E. coli* plasmid interactions in the urinary tract

Extrapolating from our results and what we know of other niches, we can estimate that F plasmids are widespread in urinary *E. coli*^42, 100, 101^. We propose a scenario that parallels how antibiotic use can select for bacteria hosting plasmids with antibiotic resistance genes^45, 102, 103^. Phage predation in the urinary tract may drive the transmission and persistence of anti-phage plasmids and, by extension, the genes linked in the plasmid, such as antibiotic resistance and virulence genes. When a urinary *E. coli* is exposed to coliphage, it can defend itself with anti-phage genes in its chromosome, plasmids, or prophage^104^. Chromosomal genes may have limits on the content that can be mutated, while prophage may require lytic activation for rapid propagation in the host population^46, 85, 105^. However, an advantage of plasmids is that they are pliable, non-essential MGEs that can be transmitted vertically and horizontally without fatally disrupting the host^106, 107^.

A major clinical issue is the increasing frequency of antibiotic resistance and virulence in bacteria, both traits that can be associated with plasmids^41, 73, 108^. Potentially, phage could drive retention and spread of clinically relevant plasmids. In this study, we presented evidence that specific urinary F plasmids can reduce permissivity to coliphage infection. These plasmids are conjugable, stable, and confer antibiotic resistances to the host. Virulence and antibiotic resistances are commonly associated with F plasmids, and predation by phage could be a very relevant selection factor in maintenance and transmission of these traits. Future studies should focus on identifying the genetic determinants in plasmids that affect phage infection. Further tests should determine to what extent urinary plasmids containing phage-protective factors can impact urinary *E. coli*, plasmid, and phage population dynamics.

## Supplemental Information

**Supplemental Table 1. Urinary *E. coli* plasmidic assembly overview**

**Supplemental Table 2. Plasmid incompatibility genes in urinary *E. coli* plasmidic assemblies**

**Supplemental Table 3. Phage spot titration results for *E. coli***

**Supplemental Table 4. Antibiotic plate growth for *E. coli* K-12 transconjugants**

**Supplemental Table 5. Presence of ORFs shared by pU0928 and pU1284 in other urinary E. coli plasmids**

**Supplemental Table 6. Phage-like sequences predicted in plasmids pU0928, pU1223, and pU1284**

**Supplemental Table 7. Predicted phage sequences in urinary *E. coli p*lasmids that have the phage-integrase ORF present in pU0928 and pU1284**

**Supplemental Figure 1. Antibiotic resistance predicted in urinary *E. coli* plasmidic assemblies**

**Supplemental Figure 2. Growth of K-12 infected with phage and stability of urinary plasmids**

**Supplemental Figure 3. Comparison of phage sequences predicted in pU0928, pU1223, and pU1284**

---

**Table 1.** Phage spot titration and antibiotic plate growth phenotype (overnight compared to 10-day passage)

|  | K-12 MG1655 | K-12 MG1655 | UMB0928 | UMB0928 | K-12 MG1655<br>*yfiQ::cm, ΔcobB* | K-12 MG1655<br>*yfiQ::cm, ΔcobB* | K-12 MG1655<br>pCA245n-cm | K-12 MG1655<br>pCA245n-cm |
| --- | --- | --- | --- | --- | --- | --- | --- | --- |
| Plasmid | None | None | pU0928 | pU0928 | pU0928 | pU0928 | pU0928 | pU0928 |
| Days passage | One | Ten | One | Ten | One | Ten | One | Ten |
| P1vir spot titration | 8 | 8 | No spot | No spot | 3 | 3 | 3 | 3 |
| Greed spot titration | 8 | 8 | No spot | No spot | 3 | 3 | 3 | 3 |
| Lust spot titration | 8 | 8 | No spot | No spot | 3 | 3 | 3 | 3 |

Note: *yfiQ* and *cobB* mutations do not affect phenotype in growth conditions utilized. Number denotes titration of phage (starting $10^9$ PFU/ml which is titration 1) at which a lysis was observed on E. coli lawn ( $\sim 10^8$ CFU/ml). A lower number indicates lysis observed only at lower titrations (i.e., higher phage PFU/ml)

|  | K-12 MG1655 | K-12 MG1655 | UMB0928 | UMB0928 | K-12 MG1655<br>*yfiQ::cm, ΔcobB* | K-12 MG1655<br>*yfiQ::cm, ΔcobB* | K-12 MG1655<br>pCA245n-cm | K-12 MG1655<br>pCA245n-cm |
| --- | --- | --- | --- | --- | --- | --- | --- | --- |
| Plasmid | None | None | pU0928 | pU0928 | pU0928 | pU0928 | pU0928 | pU0928 |
| Days passaged without plasmid selection | One | Ten | One | Ten | One | Ten | One | Ten |
| LB | + | + | + | + | + | + | + | + |
| Ampicillin |  |  | + | + | + | + | + | + |
| Chloramphenicol |  |  |  |  | + | + | + | + |
| Kanamycin |  |  |  |  |  |  |  |  |
| Spectinomycin |  |  |  |  |  |  |  |  |
| Tetracycline |  |  | + | + | + | + | + | + |

Note: “+” denotes growth in antibiotic media plate. pU0928 has a native tetracycline resistance cassette.

